# Neurotype matching in monogamous rodents is modulated by early-life sleep experience

**DOI:** 10.1101/2025.09.24.678442

**Authors:** Lezio S. Bueno-Junior, Noah E. P. Milman, Carolyn E. Jones-Tinsley, Anjesh Ghimire, Peyton T. Wickham, Yujia Hu, Bing Ye, Miranda M. Lim, Brendon O. Watson

**Author notes:** Correspondence: Lezio S. Bueno-Junior, Brendon O. Watson.

## Abstract

Studies of human sociability indicate stronger social affinity in matched-neurotype dyads (e.g., two individuals with autism or two without) compared to mixed-neurotype dyads (e.g., one individual with autism paired with one without). Is this neurotype matching phenomenon also quantifiable in non-human animals? Using deep learning tools, we analyzed dyadic male-female interactions in prairie voles, a highly social rodent species. To simulate “neurotypes”, voles were exposed to either control conditions or early-life sleep disruption (ELSD) during a critical neurodevelopmental period (postnatal days 14-21), recapitulating two features of human autism: developmental sleep disruption and later-life atypical sociability. Analogous to human studies, voles showed signs of reduced social affinity in mixed dyads compared to matched dyads, including sex-specific changes in aggression and body orientation toward the conspecific. These findings advance our understanding of social affinity, providing a framework for new studies in both animal models and humans.

## Introduction

During social interactions, we may experience feelings of closeness and affinity or, conversely, awkwardness and lack of affinity. Researchers have sought to quantify these feelings of “social chemistry” in humans, often through questionnaires to assess subjective perceptions about the social peers [1–6]. There have also been attempts to track these social properties more objectively in humans by measuring story recall, speech fluency, motion energy, facial expressions, mutual gaze, task cooperation and inter-brain synchrony [7–20].

Most of these human studies have focused on the smallest social unit – the dyad – to assess how the similarity between individuals affects social affinity, or rapport. A common strategy in these studies is to mix and match individuals with different “neurotypes”, especially those with autism spectrum disorders (ASD) or neurotypical controls, forming dyads with matched neurotypes (neurotypical-neurotypical or ASD-ASD) or mixed neurotypes (neurotypical-ASD or vice-versa). Interestingly, mixed dyads have been associated with lower affinity than matched dyads, suggesting that it is not simply the number of neurodivergent individuals in the dyad that modulates sociability [2–6,13,15,17–19]. This aligns with the growing awareness that social difficulties in neuropsychiatric spectra may partly result from interpersonal mismatches, rather than solely from individual traits [21–23].

To explore neurotype matching biologically, we can use rodent models. However, neurotype matching research has limited correspondence in the rodent model literature, despite the existence of established rodent models in social neuroscience, as well as computational ethology tools for tracking groups of animals. Rodent models include ASD-like phenotypes or genotypes (e.g., prenatal valproic acid rat model and Shank3 knockout mouse model) [24,25], which capture some components of sociability but are often limited by the focus on just one animal per assay. In turn, computational ethology tools include whole-body or body-part keypoint trackers (e.g., JAABA, idtracker.ai, DeepLabCut, SLEAP) [26–29], which capture multi-animal dynamics but fail to identify nuances, like the social role of each individual. Our group has contributed innovations to both areas. For rodent modeling, we have described that early-life sleep disruption (ELSD) impairs adult pair bonding in a monogamous rodent species: the prairie vole (*Microtus ochrogaster*), mimicking the link between developmental sleep issues and altered sociability in human ASD [30]. For computational ethology, we initially published a DeepLabCut-based system to measure body orientation and distance between two spatially-restricted prairie voles [31]. Then, more recently, we co-developed LabGym2: a postural motif tracker that identifies social roles (e.g., “chasing” versus “being chased”) in groups of freely-moving animals, including prairie vole dyads [32].

In the present study, we integrated these innovations to quantify social roles in matched versus mixed prairie vole dyads, drawing a parallel to neurotype matching studies in humans. Specifically, male and female prairie vole pups were exposed to control conditions or ELSD from postnatal days 14-21, a critical neurodevelopmental window. Upon reaching adulthood, these voles were paired into matched or mixed neurotype combinations. Behaviors were then analyzed using long-duration recordings, lasting several hours to days. We found that mixed dyads exhibit altered body orientation behaviors and higher incidence of aggressive bouts, comparable to the lower affinity found previously in mixed human dyads. Furthermore, each behavior in prairie voles varied by sex and followed a different timeline, adding both sex specificity and temporal resolution to the available knowledge from human dyad studies. These findings have implications for both the basic and clinical understanding of social affinity, in addition to offering an analytical framework for future studies in different species.

## Results

### Overview of animal pairing and behavioral analysis

We utilized two behavioral phenotypes: early-life sleep disrupted (ELSD) prairie voles and age-matched controls (Ctrl), including both sexes. This resulted in four dyad combinations: Ctrl-Ctrl, ELSD-ELSD, Ctrl-ELSD, ELSD-Ctrl (first and second acronyms represent male and female, respectively). The ELSD method was initially validated by our group [30] and is illustrated in **Fig 1A**. Briefly, home cages containing both parents and their pups – postnatal days (P) 14-21 – were placed on orbital shakers for gentle, intermittent agitation (see Methods). The pups were then weaned and housed with same-sex siblings under standard conditions until adulthood, generating cohorts of male and female ELSD voles. Ctrl voles underwent the same conditions, except for sleep disruption with the orbital shaker (**Fig 1A**). Our previous studies [30,33] have shown that ELSD reduces REM sleep time and fragments NREM sleep in prairie vole pups while preserving their body weight, corticosterone levels, and parental care. Subsequent to these acute changes, we have shown that ELSD triggers long-term changes in social affiliation (e.g., partner-directed huddling during tests for pair bond formation), cognitive flexibility, object interactions and neocortical cytoarchitecture, forming a phenotype relevant to human ASD [30,34].

**Fig 1.**
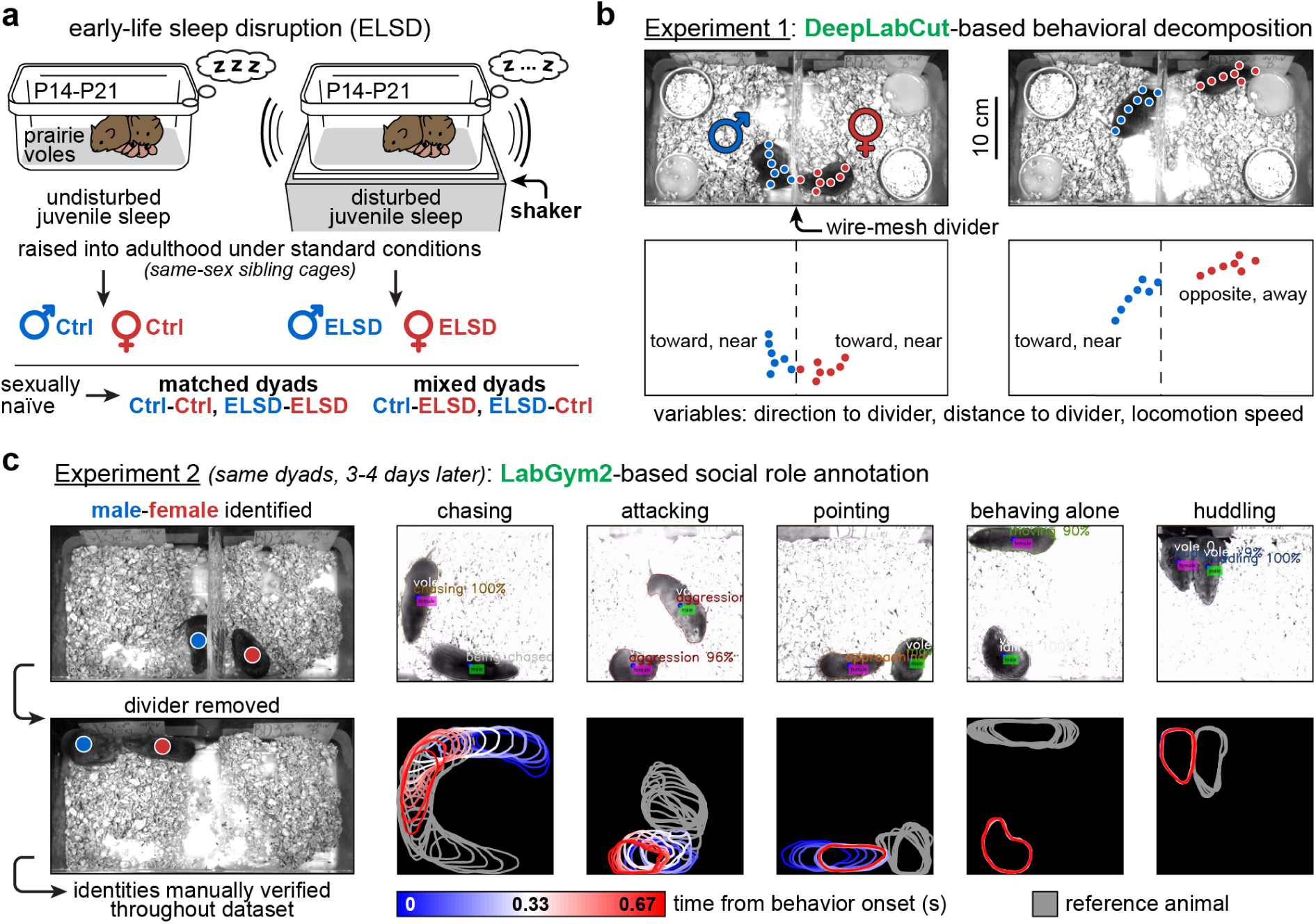
Overview of animal pairing and behavioral analysis. **A.** Early-life sleep disruption (ELSD) in prairie vole pups (post-natal days 14-21) generates phenotypes relevant to autism, as indicated by behavioral and cytoarchitectural evidence [30,34]. ELSD subjects and their controls were raised into adulthood in same-sex sibling groups under standard conditions. The adults were then assigned into matched and mixed opposite-sex dyads. **B.** Experiment 1: Male and female were filmed for 72 h while they interacted through a wire-mesh divider at the center of the cage. Cameras were positioned at an overhead angle and the animals were tracked using DeepLabCut [28], yielding cage-referenced behavioral variables. A similar method was utilized in a previous study [31] but without directly investigating dyad type. **C.** Experiment 2: After 3-4 days of single housing in the vivarium, the same dyads were placed back together into the recording setup, this time for 4 h and without the divider. This resulted in a different kind of dataset comprising partner-referenced social role categories, annotated using LabGym2 [32] and manual validation (see Methods).

Here, ESLD effects on dyadic behavior are explored more deeply thanks to computational ethology. We describe two strategies. <u>Experiment 1</u>: body-part keypoint tracking (DeepLabCut) [28] of mutually naïve male-female pairs separated by a wire mesh divider (**Fig 1B**). <u>Experiment 2</u>: whole-body behavioral classification (LabGym2) [32] of the same male-female pairs, but without the divider (**Fig 1C**).

<u>Experiment 1: initial divided cohabitation</u> (**Fig 1B**; **S1 Movie**). This experiment consisted of 72-h overhead video recordings where voles interacted through a wire-mesh divider at the center of the cage. This was the first encounter these animals had with each other. While this setup imposed a physical limitation on social interactions, it yielded uniform postural and spatial measures throughout recordings. This allowed us to decompose these measures into three DeepLabCut-derived variables: body direction to divider, distance to divider and locomotion speed (**Fig 1B**). In a previous study [31], we characterized the sex-specific impacts of ELSD to these variables, including alterations in male body orientation and female circadian activity. However, we did not examine neurotype matching effects in that study; such effects are reported here.

<u>Experiment 2: subsequent undivided cohabitation</u> (**Fig 1C**; **S2 Movie**). This experiment consisted of 4-h overhead video recordings using the same voles, three to four days after Experiment 1, meaning that the opposite-sex mates had already been exposed to each other at the time of Experiment 2, i.e., dyad assignments were preserved between experiments (see Methods). In this setup, the animals were allowed to engage in freely moving partner-referenced behaviors without a divider. This resulted in an enriched behavioral repertoire, which is more difficult to analyze with DeepLabCut but amenable to newer LabGym2-based behavioral classification (**Fig 1C**). One novel aspect of LabGym2 – a software that we co-developed [32] – is the ability to categorize the social role of each individual (e.g., “chasing” versus “being chased”). We utilized this capability in combination with manual validation to ensure reliable identity tracking and sex specification per behavior. This resulted in the labeling of five behavior categories: *chasing*, *attacking*, *pointing*, *behaving alone*, and *huddling* (**Fig 1C**). Here, “*pointing*” combines sniffing behaviors with a freezing-like behavior that we often observed after each aggressive bout (see **S2 Movie**). See Methods for details.

Therefore, our two experiments yielded very different analyses. Nevertheless, neurotype matching effects emerged in both experiments, as reported below.

### Experiment 1: Mixed dyads showed exacerbated sex differences in body orientation

In Experiment 1, male and female interacted through a wire-mesh divider. **Fig 2** describes these interactions over 72 h (1-h bins, see x-axes), with graph columns displaying DeepLabCut-derived behavioral decomposition variables. **Fig 2A** starts by comparing sexes irrespective of dyad grouping. When analyzing direction to divider, we found that male voles spent significantly more time facing toward the divider than females throughout the recording. For distance to divider, both sexes preferred areas near the divider (4-6 cm) during the initial 12 h of recording; this variable later plateaued at around 6 cm (halfway between divider and opposite cage wall), suggesting habituation. For locomotion speed, both sexes exhibited higher locomotion in the initial 12 h of recording; this variable later stabilized to lower basal levels, also suggesting habituation (**Fig 2A**). These results corroborate our previous findings [31] that sex-specific body orientation patterns persist even after voles habituate to the environment, as indicated by the stabilization of locomotion and area preference after the initial 12 h. See **Table 1** for statistics and **S1 Movie** for representative behaviors.

**Fig 2.**
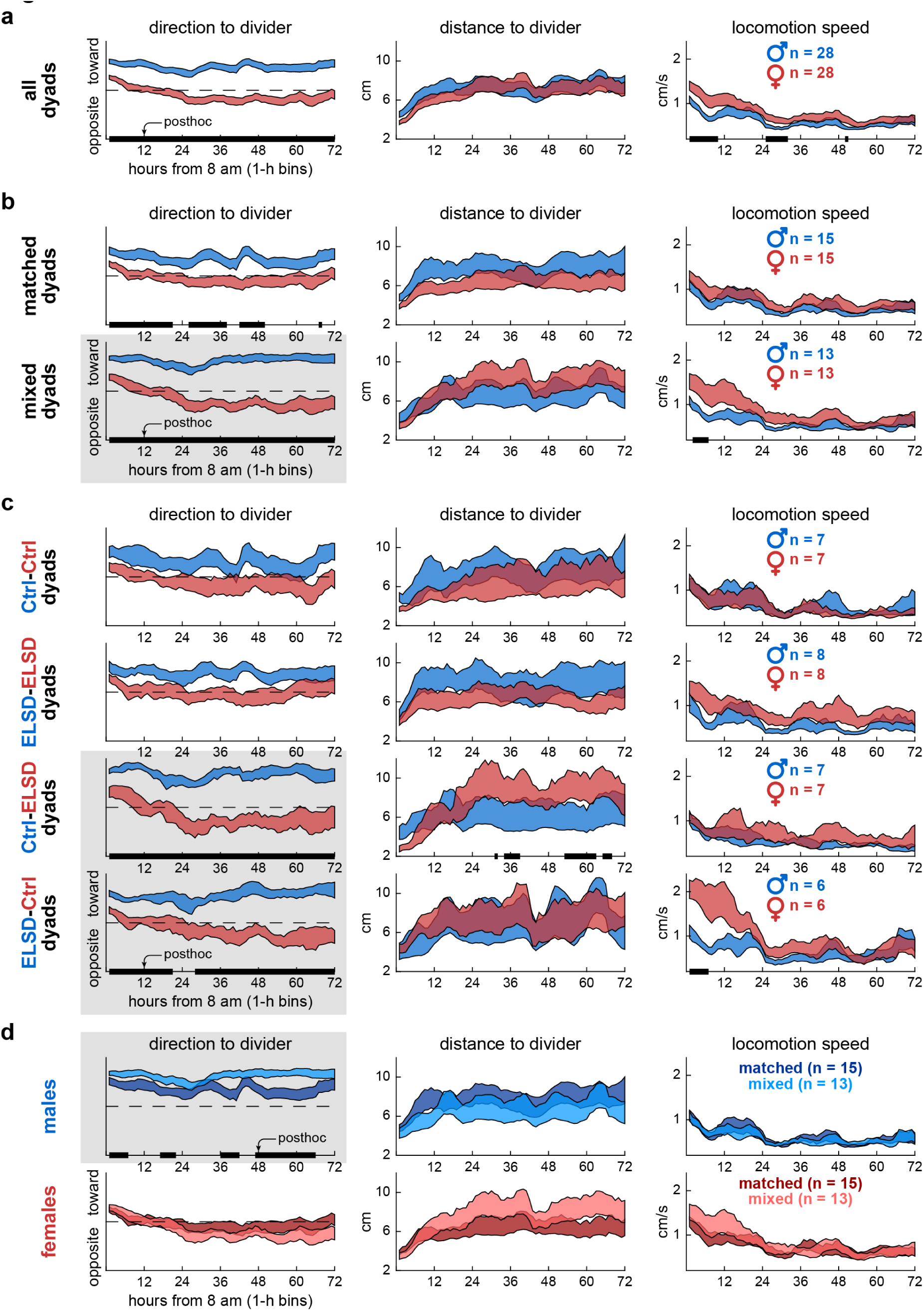
Experiment 1: Mixed dyads showed exacerbated sex differences in body orientation. **A.** Time series data (72 h, 1-h bins) showing sex differences in direction to divider across all dyads. Males spent more time toward the divider than females throughout the recordings. The curves represent mean +/-standard error aggregated across animals, regardless of individual dyad assignments. The black bars on the x-axes represent posthoc differences, after two-way ANOVA with time bins as repeated measures. **B.** Sex differences in direction to divider were more pronounced in mixed dyads (highlighted with gray background). Distance to divider and locomotion speed were less sensitive to dyad type, illustrating that behavioral decomposition was important to capture the effects shown here. **C.** Further specification into the four dyad subtypes. Sex differences in direction to divider were confined to the mixed-dyad subtypes (Ctrl-ELSD, ELSD-Ctrl). Additional smaller effects were observed in distance to divider and locomotion speed, specific to each mixed-dyad subtype. **D.** Same groupings as panel B but comparing matched and mixed dyads within each sex. The tendency to behave toward the divider was higher in males and further increased in mixed-dyad males. See **Table 1** for statistics and **S1 Movie**.

By parsing these sex differences into dyad types, we begin to reveal the novel findings of this study. **Fig 2B** shows that while sex differences in direction to divider were significant in both matched and mixed dyads, these differences were more pronounced in mixed dyads and persisted for 72 h (**Table 1**). **Fig 2C** further divides male and female voles into the four juvenile sleep status combinations. In this case, sex differences in direction to divider were significant only in mixed dyads (Ctrl-ELSD, ELSD-Ctrl). Matched dyads, including Ctrl-Ctrl and ELSD-ELSD, showed no significant sex differences (**Table 1**). These results indicate that body orientation behaviors did not change linearly with the number of ELSD individuals in the dyad; rather, neurotype matching was the main factor modulating body orientation.

Additional sex differences were specific to mixed dyads and confined to more localized time periods. We observed that ELSD females in Ctrl-ELSD dyads maintained a greater distance from the divider more than 36 hours into the session, while Ctrl females in ELSD-Ctrl dyads exhibited higher locomotion speed during the initial 12 hours compared to males (**Fig 2C**). However, these sex differences were modest compared to the prominent results in direction to divider (**Table 1**). **Fig 2D** further highlights direction to divider as a sensitive variable, this time comparing matched and mixed dyads within each sex. Males spent significantly more time toward the divider in mixed compared to matched dyads, indicating heightened female-directed orientation in mixed-dyad males (**Table 1**).

Overall, **Fig 2** shows that a supra-individual factor (neurotype matching) preferentially modulated one behavioral component (body direction) in one sex (males). Next, we describe that neurotype matching remained an important factor in a different environment.

### Experiment 2: Mixed dyads showed higher female-to-male aggression

In Experiment 2, the wire mesh divider was not present, and animals were able to interact in an unrestricted fashion. **Fig 3** is structured similarly to the results above, with two differences: the recordings lasted 4 h (3-min bins, see x-axes) and the graph columns show LabGym2-based behavior categories. We again start by comparing sexes irrespective of dyad grouping in **Fig 3A**. The social roles of *chasing* and *attacking* were significantly more prevalent in female voles, especially in the first hour of recording (insets), after which these variables decayed to lower basal levels. The social role of *pointing*, which included sniffing and other behaviors (see Methods), was evenly distributed across sexes and showed a more gradual decline over the 4-h period (see lower F values in effects of time in **Table 2**). The role-unspecific categories of *behaving alone* and *huddling* (i.e., behaviors that considered the full dyad, see Methods) also showed gradual time courses, but in opposite directions: *behaving alone* decreased, while *huddling* proportionally increased over time (**Fig 3A**). In addition, *behaving alone* and *huddling* were the most frequent behaviors across **Fig 3**, as indicated by the probability axes. See **Table 2** for statistics.

**Fig 3.**
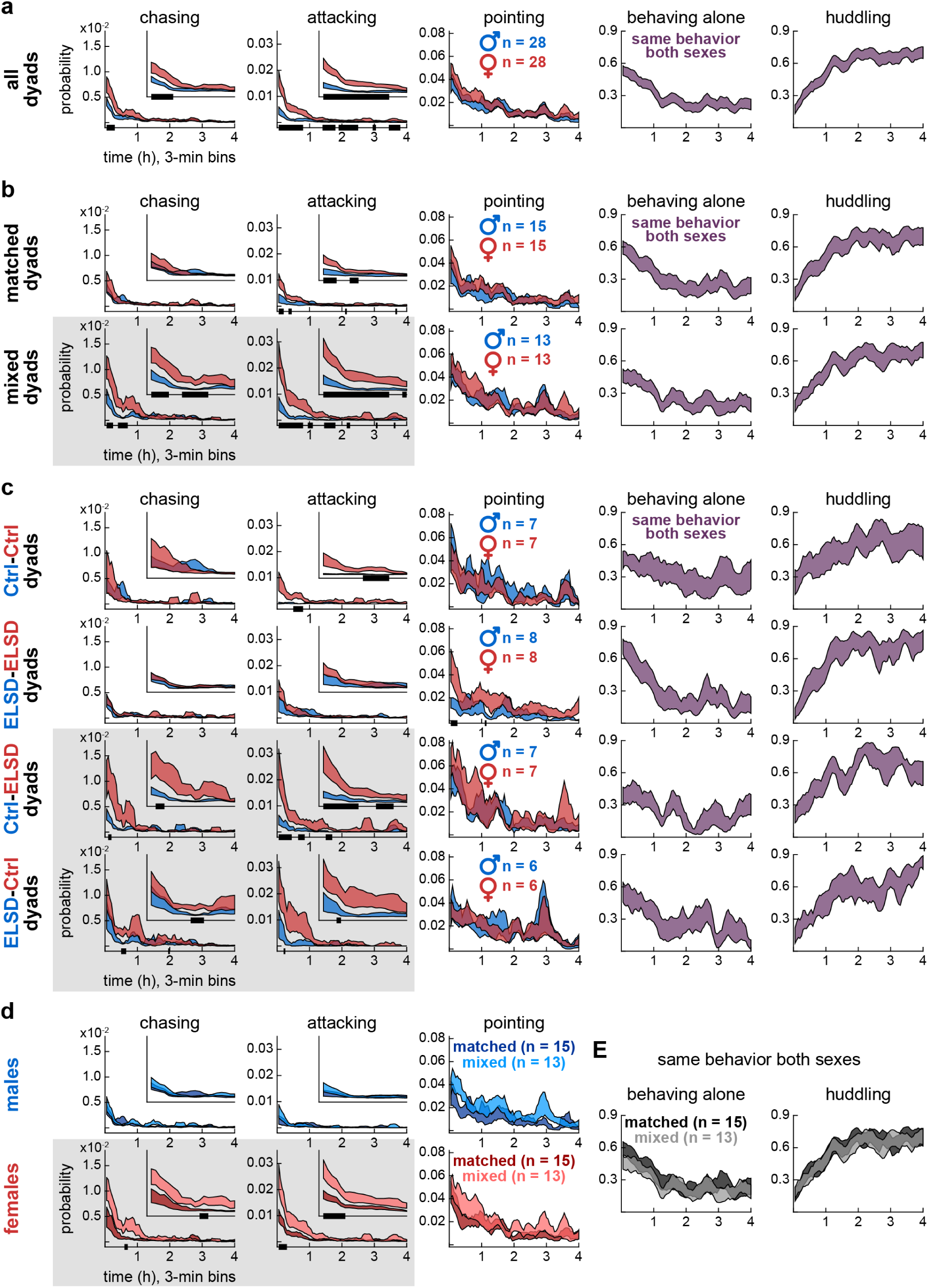
Experiment 2: Mixed dyads showed higher female-to-male aggression. **A.** Time series data (4 h, 3-min bins) showing sex differences in the likelihood of chasing and attacking across all dyads. Females were more likely to chase and attack than males, especially in the initial hour of recording. The curves represent mean +/- standard error aggregated across animals, regardless of individual dyad assignments. The black bars on the x-axes represent posthoc differences, after two-way ANOVA with time bins as repeated measures. Inset plots show probability within the first hour. **B.** Sex differences in the likelihood of chasing and attacking were more pronounced in mixed dyads (highlighted by gray background). Pointing behaviors, including face-to-face, face-to-body and anogenital sniffing, showed no sex or dyad type specificity, making them useful comparators (see Methods for details). The role-unspecific categories of alone and huddling were more dominant and mutually exclusive, illustrating the transition from high activity (periods of moving and idling alone punctuated by chasing, aggression and sniffing bouts) to low activity (periods of continuous huddling). **C.** Further specification into the four dyad subtypes. Sex differences in the likelihood of chasing and attacking were stronger in the mixed-dyad subtypes (Ctrl-ELSD, ELSD-Ctrl). **D.** Same groupings as panel B but comparing matched and mixed dyads within each sex. Mixed-dyad females were more likely to chase and attack than matched-dyad females. See **Table 2** for statistics and **S2 Movie**.

Despite their lower occurrence rate, *chasing* and *attacking* are the behaviors with the greatest differential trajectories across types of dyads. As shown in **Fig 3B**, female voles were more likely to chase and attack the males in mixed than matched dyads (**Table 2**). Consistently, in dyad subgroups (**Fig 3C**), female-to-male *chasing* and *attacking* were more frequent in both Ctrl-ELSD and ELSD-Ctrl dyads (**Table 2**). Matched ELSD-ELSD dyads showed no sex differences in *chasing* and a comparatively small difference in *attacking* (**Table 2**), reinforcing that these behaviors did not change linearly with the number of ELSD individuals in the dyad, similar to the findings in dyads behaving across a divider. Additional sex differences were observed in *attacking* (Ctrl-Ctrl) and *pointing* (ELSD-ELSD) (**Fig 3C**), but these were also modest compared to the sex differences in *chasing* and *attacking* behaviors found in mixed dyads (**Table 2**). Furthermore, **Fig 3D** shows within-sex comparisons between dyad types, confirming that mixed-dyad females were more likely to chase and attack than matched-dyad females. No dyad type differences were observed in the other behaviors, i.e., *pointing*, *behaving alone* and *huddling* (**Fig 3D-E**; **Table 2**), again highlighting female-to-male *chasing* and *attacking* as the most sensitive behaviors in this analysis.

Therefore, neurotype matching preferentially modulated female-to-male aggression in the undivided cage. Below we link Experiments 1 and 2 via correlations within dyads.

### Linking divided cohabitation to undivided cohabitation: Neurotype matching affected correlations among behaviors

The same dyad assignments were used in Experiment 1 (first male-female encounter and 72-h divided cohabitation) and Experiment 2 (second encounter and 4-h undivided cohabitation), with an interval of three to four days between experiments. This gave us the opportunity to quantitatively link these experiments. Thus, we averaged all behavioral variables per animal (three variables from Experiment 1, five from Experiment 2) and explored linear correlations among such variables. **Fig 4A-B** illustrates correlations between direction to divider in Experiment 1 and all behavioral categories of Experiment 2. Each data point in **Fig 4A-B** is an individual animal. Separate analyses were performed per sex and major dyad type (see Methods).

**Fig 4.**
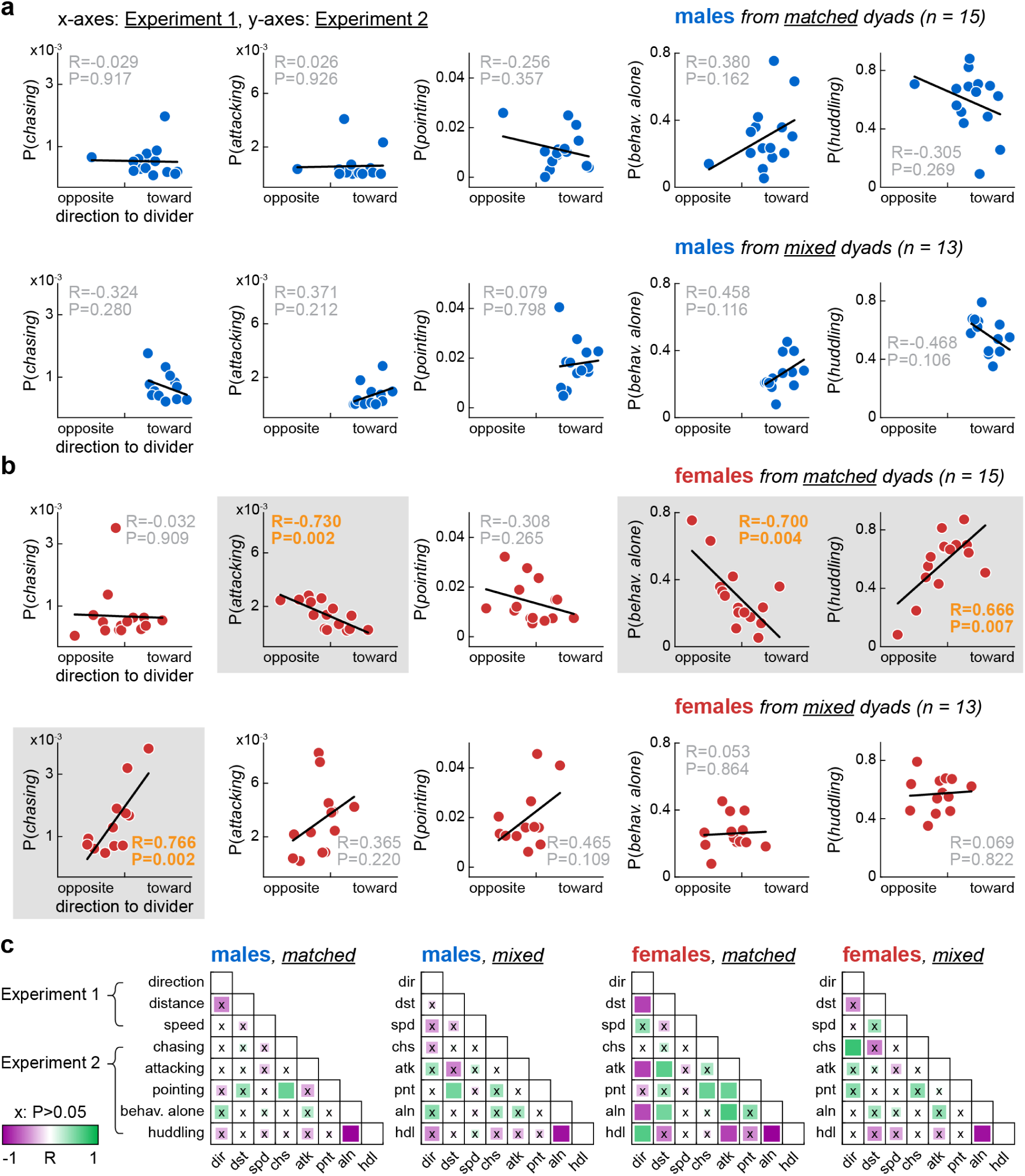
Linking divided cohabitation to undivided cohabitation: Neurotype matching affected correlations among behaviors. **A.** Correlations between the most sensitive variable from Experiment 1 (72-h divided cohabitation), direction to divider (x-axes), and behavior probabilities from Experiment 2 (4-h undivided cohabitation; y-axes), averaged per male individual (data points). **B.** Same but for females. Significant correlations revealed further distinctions between dyad types, highlighted by gray backgrounds. Matched-dyad females formed an affiliation-driven spectrum: a higher tendency to behave toward the divider in Experiment 1 correlated with higher sociability 3-4 days later, in Experiment 2 (lower likelihood to attack and behave alone, higher likelihood to huddle). This pattern was absent in mixed-dyad females; instead, they exhibited a positive correlation between direction to divider and chasing behavior. **C.** Comprehensive view of all correlation pairs. R and P values are coded by color and square size, respectively. Note that the scatter plots in panels A-B correspond to the leftmost column of each correlation matrix.

Significant correlations were restricted to female voles and differed between matched versus mixed dyads (**Fig 4B**; see plots with gray background). *Chasing*: positively correlated with direction to divider in mixed. *Attacking*: negatively correlated with direction to divider in matched. *Behaving alone*: negatively correlated with direction to divider in matched. *Huddling*: positively correlated with direction to divider in matched (**Fig 4B**). Thus, a female that oriented more toward the divider in Experiment 1 was more likely to engage in different behaviors in Experiment 2 depending on the dyad type: affiliative behaviors in matched, chasing behaviors in mixed. This suggests that, depending on the dyad type, the same behavior in Experiment 1 (body orientation) may have been driven by a different underlying state. **Fig 4C** presents correlation matrices for all variable pairs, providing a comprehensive view of both sex and dyad type differences. The leftmost columns in each matrix of **Fig 4C** (direction to divider) contain the correlations emphasized above. Other correlations were found within each experiment, including negative correlations between alone and huddling, consistent with their opposing trends in Experiment 2 (**Fig 3**).

### Female-to-male behavior transitions differed between matched and mixed dyads

The analyses above ignore within-dyad actions and reactions at finer temporal scales. How likely is behavior B in vole 2 after behavior A in vole 1? Do these likelihoods change with dyad type and/or sex directionality (male-to-female, female-to-male)? In **Fig 5** we explore these questions using behavior transition probabilities, thus capturing temporal sequence patterns from each dyad type and sex directionality. Data from the video frames were divided into 1-s time bins; this temporal resolution was deemed optimal to capture all transitions between behaviors, including to and from aggression bouts – the quickest behaviors in our dataset (**S2 Movie**). We then conducted this analysis separately per major dyad type (matched, mixed) and sex directionality (male-to-female, female-to-male).

**Fig 5.**
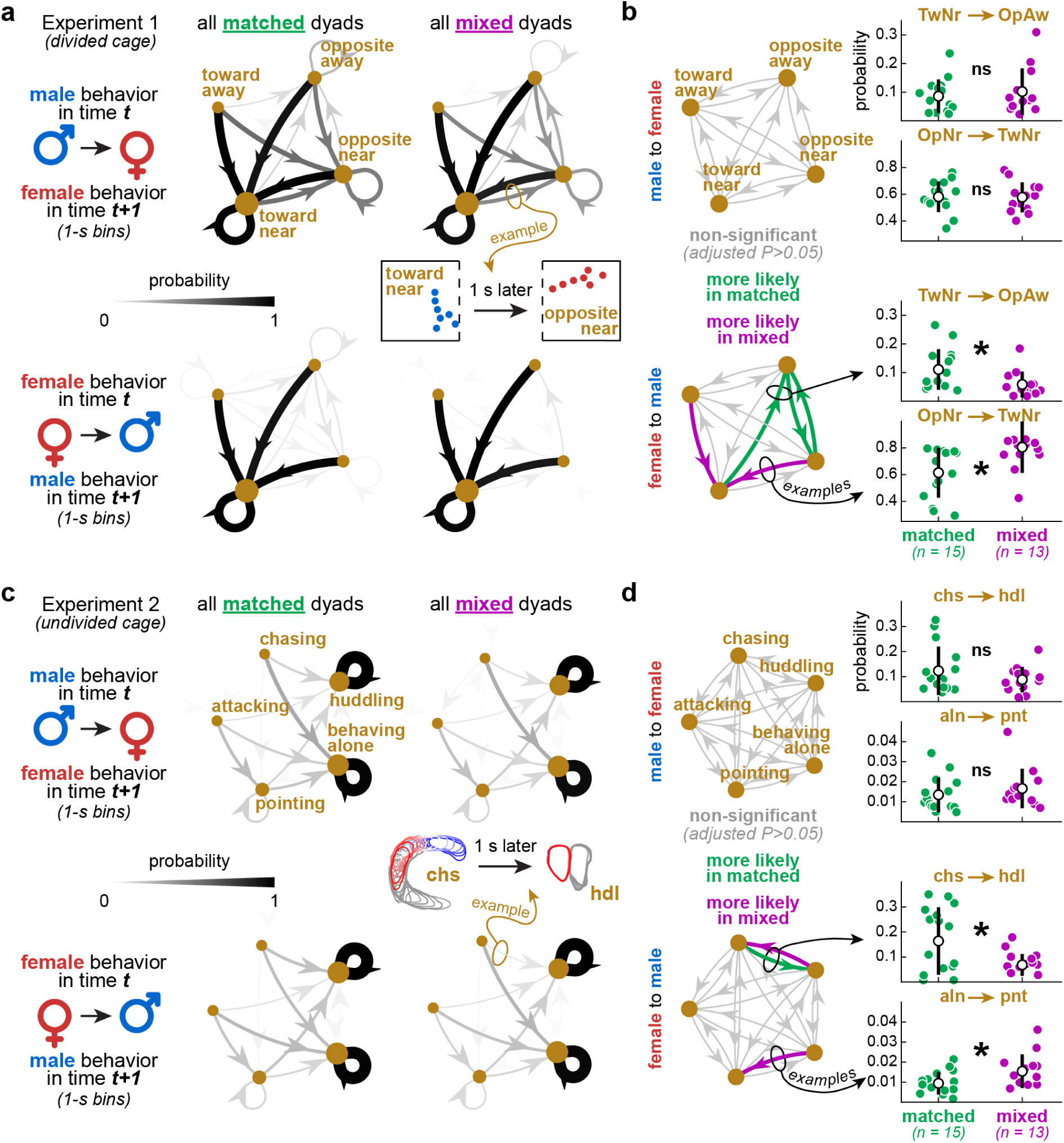
Female-to-male behavior transitions differed between matched and mixed dyads. **A.** Data from Experiment 1 (divided cohabitation). Direction and distance to divider were time-binned (1-s), categorized (toward-near, opposite-near, toward-away, opposite-away), and analyzed as male-to-female and female-to-male transition probability matrices (see Methods). Directed graphs (averaged across dyads) illustrate patterns that varied with both dyad type and sex directionality. **B.** Quantitative comparisons between dyad types within each sex directionality, using one transition matrix per dyad, rather than the averages in panel A. Behavior pairs were used as repeated measures in this analysis (see Methods). Self-loops were excluded to focus on inter-behavior relationships. The directed graphs highlight which transitions differed statistically between dyad types. The scatter plots exemplify the distributions underlying each transition edge (each data point is a dyad). Mean +/- standard deviation bars accompany the scatter plots. **C-D.** Same as panels A-B but from Experiment 2 (undivided cohabitation). Overall, these patterns suggest that matched dyads showed more variation in body re-orientation behaviors between the females and males, as well as a higher likelihood of female chasing leading to huddling, among other interpretations. See Results and Discussion. See **Table 3** for statistics.

**Fig 5A-B** shows data from Experiment 1 (divided cohabitation). The most informative variables in that dataset – direction and distance to divider – were continuous and therefore required discretization prior to transition analysis. Thus, we discretized direction to divider into “toward” or “opposite”, and distance to divider into “near” or “away”, resulting in four categories: *toward-near*, *opposite-near*, *toward-away*, *opposite-away* (see Methods). The directed graphs in **Fig 5A** illustrate transitions among these categories, averaged across dyads. The edges represent probabilities, coded by line width and grayscale (**Fig 5A**). For this analysis, we focused on the initial 12 h of the 72-h recording, when the animals were more active (**Fig 2**).

The main visual pattern in all graphs is the convergence to, and recurrence of, *toward-near* (black edges): these represent the highest transition probabilities in both sex directionalities (**Fig 5A**). Particularly in the male-to-female graphs, the convergence to, and recurrence of, *opposite-near* (dark gray edges) also showed relatively high probabilities. All other transitions were less likely (light gray edges) (**Fig 5A**). The more distributed patterns in the male-to-female graphs indicate higher variation: female voles were more active and rotated themselves more (**Fig 2**, **S1 Movie**), thus increasing the chances of male behaviors being followed by both body orientation categories in the females. In contrast, the female-to-male graphs indicate lower variation: all female behaviors were more likely followed by *toward-near* in the males, simply because this behavior was predominant in males (**S1 Movie**).

Of interest to this study, we found differences between matched and mixed dyads, specifically in female-to-male transitions. For quantification, transition probability matrices were calculated per dyad, reshaped into dyad x edge matrices (edges were used as repeated measures), and compared between matched and mixed dyads, resulting in **Fig 5B** (self-loops were excluded; see Methods). In mixed dyads, two female-to-male transitions were more likely: *opposite-near* to *toward-near* and *toward-away* to *toward-near* (**Fig 5B**). We interpret that this was driven by the tendency of mixed-dyad males to spend more time toward the divider, as reported earlier (**Fig 2**). In matched dyads, three female-to-male transitions were more likely: *toward-near* to *opposite-away*, *opposite-away* to *opposite-near*, and vice-versa for the latter (**Fig 5B**). These re-orientation patterns between male and female were infrequent (see light gray edges in **Fig 5A**) but could be speculated as body posture signals. **Fig 5B** additionally contains examples of the distributions behind each behavior pair, shown here as mean +/- standard deviations (each data point represents a dyad). See **Table 3** for statistics.

**Fig 5C-D** shows the same analysis, but from the full 4 h of Experiment 2 (undivided cohabitation), with five labeled behaviors during free interaction. As illustrated by the average directed graphs in **Fig 5C**, the highest transition probabilities were found in the recurrence of *behaving alone* and *huddling* (black self-loops). This was expected, as *behaving alone* and *huddling* were both prevalent and self-sustaining, as shown previously (**Fig 3**). For quantification, we excluded such loops to focus on the behavior pairs, as described above, resulting in **Fig 5D**. In mixed dyads, two female-to-male transitions were more likely: *huddling-to-chasing* to *alone-to-pointing* (**Fig 5D**). This could reflect higher activity overall in mixed dyads, including shorter and more intermittent periods of huddling and more frequent alone-to-pointing re-encounters. However, these transitions showed very low probabilities (hardly visible in **Fig 5C**; see also y-axes in **Fig 5D**) and thus we interpret them cautiously. A more robust observation was that female-to-male *chasing-to-huddling* was more likely in matched dyads (**Fig 5D**). Thus, female chasing in matched dyads was possibly more affiliative, or huddling-seeking (**S2 Movie**). This suggests the existence of different states underlying the same behaviors depending on dyad type, echoing an interpretation we proposed in the previous subsection.

Therefore, aside from the specificities of each behavior pair, **Fig 5** shows that neurotype matching influences actions and reactions on the scale of seconds, reinforcing the time series and correlation analyses reported earlier.

## Discussion

We utilized algorithms for annotation and decomposition of social behaviors to examine neurotype matching in rodents. Opposite-sex prairie voles were assigned to either matched or mixed-neurotype dyads, resembling human studies involving ASD. The voles were then filmed in two experiments: initial divided and subsequent undivided cohabitation recordings. Dyad type effects emerged from both experiments in a correlated manner, suggesting robust underlying mechanisms. Our observations varied by sex and spanned multiple temporal scales, including body orientation patterns over days, incidence of aggression over hours, and behavioral transitions over seconds. These findings might influence further development of behavioral paradigms and studies integrating social behavior and physiology.

### Juvenile sleep and atypical sociability

Neurotype matching studies in humans often involve mixing and matching neurotypical and neurodivergent individuals, especially those with ASD [2–6,13,15,17–19]. Here, “neurodivergent” corresponds to adult prairie voles exposed to ELSD during post-natal days 14-21, a method described previously by our group [30,33]. ELSD causes fragmentation of non-rapid eye movement sleep (NREM) and reduces both duration and gamma power of rapid eye movement sleep (REM), as recorded electrographically from prairie vole pups. Body weight, corticosterone levels, and parental care are all unaffected by ELSD, indicating that this method selectively targets juvenile sleep physiology [30,33]. Despite this selectivity, ELSD provokes long-lasting behavioral and neurobiological alterations into adulthood. Behavioral alterations include impaired cued fear extinction and reduced male-to-female partner preference expression. Neurobiological alterations include cytoarchitectural markers of excitatory-inhibitory imbalance in brain regions involved in sensory and cognitive processing, such as increased expression of inhibitory neurons in somatosensory cortex, as well as increased dendritic spine density and decreased presynaptic glutamate labeling in prefrontal cortex [30,34]. Similar neurobiological changes are present in human ASD, making prairie vole ELSD relevant for investigating ASD and other disorders linked to poor juvenile sleep [35].

As reviewed previously [35–37], sleep disruption during infancy has been associated with ASD in humans. Brain maturation processes in humans, such as axonal myelination and synaptic pruning, depend on healthy sleep during development. Disrupted sleep may interfere with these processes and increase vulnerability to ASD-like symptoms, including social difficulties [35–37]. Since neurotype matching modulates behavior in both prairie vole ELSD (present study) and human ASD [2–6,13,15,17–19], we can speculate that some of these social properties may trace back to sleep quality during development. This is, of course, a more complex question. First, the relationship between sleep disruption and ASD is recognized as a bidirectional and multifaceted comorbidity, with sleep disturbances both amplifying and being amplified by ASD [36,37]. Second, human ASD is linked with other factors not directly related to sleep, such as heritability and perinatal exposure to toxicants [38]. And third, juvenile sleep is sensitive to circumstances not directly related to ASD, such as respiratory problems and household conditions [39,40]. All these factors can lead to (and interact with) ASD [41], making it difficult to isolate the role of juvenile sleep in ASD, let alone dyad type properties. Still, our results suggest that juvenile sleep does play a role in these social properties – a role that we found to be discernible in non-human animals, despite the elusive mechanisms. Future studies could disentangle poor juvenile sleep from other environmental and genetic factors using different rodent models, furthering the study of non-human neurotype matching and complementing human ASD research.

### Markers of social affinity across neuropsychiatric spectra

Human ASD encompasses a myriad of features, including subtle presentations of ASD, which can be challenging to detect and often go undiagnosed [42,43]. Likewise, ELSD prairie voles do not show overt behavioral changes. In our experience, these changes can only be detected with quantification, whether using manual [30] or computerized [31] tracking. Thus, ELSD prairie voles might represent a nuanced ASD-like spectrum, analogous to nuanced neuropsychiatric spectra in humans. Our experiments add complexity to this question. Despite the elusive nature of these spectra, their phenotypes can still supply us with controllable “reagents” to manipulate “social chemistry”. The “purity” of these reagents (i.e., phenotypes) certainly vary among species or between laboratory and naturalistic settings, yet they still seem to bring about some form of chemistry, as suggested by the literature on human dyads and our study in prairie voles. Furthermore, dyadic behavior in humans has been investigated beyond ASD, including studies on dysphoria, attention deficit disorder, and decision making [1,7–9], all of which reinforcing that mixing or matching different phenotypes (or social styles) can modulate behavioral performance and sociability. This indicates that social matching properties are measurable across species and phenotypes, with implications for basic science, mental health, and a better understanding of reciprocity and empathy [21–23].

These questions warrant further investigation using data-intensive metrics of social affinity. Recent human ASD studies involving matched and mixed dyads used objective metrics from video, audio and electroencephalography, including motion energy, mutual gaze, backchanneling, verbosity, smile synchrony, and inter-brain synchrony [12,13,15,17,18]. Generally, these studies also support neurotype matching properties, though some metrics are inconsistent with this trend. Such inconsistency might be attributed to low temporal resolution, as these studies used aggregated metrics across sessions when comparing dyad types. Time series analyses, like those here, could be employed in future studies to extract detailed information from each metric, potentially identifying optimal metrics for real-time sociability tracking and translatability across species. Other human dyad studies, unrelated to ASD or neurotype matching, used additional metrics, including task cooperation, motor coordination, smell-induced arousal, and functional brain imaging or spectroscopy [14,20,44–49]. This variety of methods suggests promising avenues for studying dyadic behavior across species and neuropsychiatric disorders.

### Sex differences and ethological nuance

In our male-female grouping design (Ctrl-Ctrl, ELSD-ELSD, Ctrl-ELSD, ELSD-Ctrl) we considered both sex and dyad type. This revealed dyad type effects both between and within sexes in prairie voles. Between-sex and within-sex interactions have also been recognized in human ASD. Core ASD traits seem to vary equivalently between sexes (e.g., communication impairments, repetitive behaviors), whereas associated ASD traits seem to vary independently within each sex (e.g., hyperactivity in males, depression in females) [50–52]. To our knowledge, these sex-related variables have not been experimentally manipulated in studies on neurotype matching and ASD. As shown here with prairie voles, sex differences in body direction (males spent more time toward the divider) and aggression (females chased and attacked more) were exacerbated in mixed dyads. We can now speculate that neurotype matching could also amplify sex differences in humans and other species. Investigating these questions could provide deeper insight into sex differences in human ASD or ASD-like models.

Moreover, we found that female body orientation in divided cohabitation (Experiment 1) led to different outcomes in undivided cohabitation (Experiment 2) depending on the dyad type: increased affiliation in matched dyads (less attacking, more huddling) and decreased affiliation in mixed dyads (more chasing, less chasing-to-huddling transitions). Therefore, a female behavior that looked the same – body direction – may have been driven by different underlying states depending on the dyad type, e.g., female body direction was possibly more huddling-oriented in matched dyads. Interestingly, these states in females may have influenced (and been influenced by) the males. In mixed dyads, males spent more time toward the divider in Experiment 1, and once the divider was removed in Experiment 2 the females attacked more frequently. Thus, females may have reacted proportionally to the males’ approaches upon removal of the divider, which could be another explanation why aggression was more frequent in mixed dyads. Aggressivity, affiliation and other behavior types (e.g., courting, foraging) are indeed recognized to vary with sex, social hierarchy and environmental context [53–55], suggesting an interplay between internal and expressed states. We propose that matching and mixing rodent “neurotypes” like we did here is a promising approach to investigate these states.

### Concluding remarks

This study reveals that neurotype matching is a quantifiable phenomenon in rodents. As computational ethology continues to optimize social behavior tracking, the emergent properties of social groups may become more tangible. This progress may reshape behavioral paradigms, from dyad-based studies to freely moving partner preference studies, strengthening the link between animal models and human sociability.

## Materials and methods

### Subjects

Prairie voles (n = 28 each sex) were sourced from a genetically diverse pool of 13 breeders in an accredited, fully-contained laboratory facility (VA Portland Health Care System). Litters containing both male and female sibling pups (2-9 pups each breeder pair) were raised with their parents (no cross fostering) until post-natal day 21 (P21). During the P14-21 period, specifically, the litters were subjected to either control (Ctrl) or early-life sleep disruption (ELSD) conditions (details below). At P21, pups were weaned into same-sex sibling groups and remained at the same vivarium until approximately P90. The sibling groups were then shipped to the University of Michigan Medical School for the primary recordings of this study.

Consistent housing conditions were maintained across both institutions, including 14:10 h light/dark cycle (lights on at 5:00 am), temperature (20-23 °C), ventilation (double-filtered outside air, negative pressure), humidity (30-70%), bedding (rolled paper pellets), environmental enrichment (cotton nestlets and wooden blocks/sticks) and *ad libitum* water and food (rabbit chow, corn, cracked oats). Each week, the animals were transferred to a new cage with fresh bedding and enrichment. All procedures in this study were approved by the Institutional Animal Care and Use Committees (IACUC) of VA Portland Health Care System (protocols 6069 and 3652) and University of Michigan Medical School (protocol 00011423).

Our animals originated from a colony at Emory University, itself derived from wild-captured animals in the State of Illinois, USA. To preserve genetic diversity, bi-annual exchanges of animals occur among several institutions, including North Carolina State University, University of California Davis, University of Colorado Boulder, and Florida State University.

### Early-life sleep disruption

Prairie vole ELSD phenotypes of both sexes were generated by placing home cages with P14-21 pups and their parents on an orbital shaker for intermittent gentle agitation (10 seconds on, 100 seconds off, 110 rotations per minute). ELSD animals were provided hydrogel instead of water bottles to prevent water spillage during agitation. Ctrl animals experienced the same conditions except for agitation (**Fig 1A**). As previously described by our group [30,33,34], ELSD disturbs sleep in prairie vole pups without affecting their hormonal markers of stress, or the parental care they receive. The acute effects of ELSD are, therefore, confined to sleep electroencephalography (EEG) parameters, including fragmentation of NREM sleep and reduction of REM sleep time by more than 20% over a 24-hour period [30]. Given this support from our previous studies, here we used identical cage agitation parameters without EEG validation, resulting in a non-invasive (purely behavioral) experimentation timeline.

### Dyad assignment and age at recording

Following the ELSD or Ctrl procedures at P21, the pups were weaned into same-sex groups of 2-4 siblings per cage until adulthood, resulting in four types of sibling groups (Male-Ctrl, Male-ELSD, Female-Ctrl, Female-ELSD). Sexually naïve opposite-sex adults were then randomly assigned into matched dyads [Male-Ctrl/Female-Ctrl (n=7), Male-ELSD/Female-ELSD (n=8)] or mixed dyads [Male-Ctrl/Female-ELSD (n=7), Male-ELSD/Female-Ctrl (n=6)] prior to recordings, for a total of 28 dyads (**Fig 1A**).

Our recording setup could accommodate up to four dyads at a time (details below). Since there were 28 dyads in total, the recordings were divided into seven cohorts of four dyads. Furthermore, an interval of 2-3 weeks was necessary between cohorts due to the duration of the experiments combined with preparatory activities, like cage cleaning and file management. This resulted in a four-month period to record all 28 dyads. Thus, the ages of the animals at the time of recording varied with a mean +/- standard error of 185.8 +/- 6.5 days of age. However, the individuals of each dyad were still age-matched, with an age difference of 11.1 +/- 2.2 days, randomly distributed across dyad types.

### Recording setup

For recording, we developed a vibration-free optomechanical assembly using metal parts from a vendor (ThorLabs), as described previously [31]. This setup holds two infrared-sensitive grayscale cameras (Basler, acA1300-60gm) positioned at 90° overhead angle. Each camera was mounted to a fixed focal length lens (Edmund Optics, 6 mm UC Series) and surrounded by four infrared LED illuminators. Each camera/lens was able to frame two home cages, hence the ability to record four dyads at a time. This system was installed in a room with light/dark phases, which however did not affect imaging brightness, as it relied on infrared reflectance. The cameras were connected to a computer via a network adapter (Intel Pro 1000/PT), and data were stored on a redundant array of independent disks (RAID). Recording settings were configured via Pylon software (Basler) with the following parameters: 8-bit grayscale depth, frame width x height of 800 x 896 pixels, no binning, 1328 kbps data rate, 20 Hz frame rate. Exposure and brightness were adjusted prior to each recording by manipulating the lenses and infrared illuminators, without digital adjustments. Data were acquired into mp4 files (H.264 coded) using StreamPix software (NorPix). For the 72-h recordings of Experiment 1, specifically, recordings were acquired into 6-h mp4 files to ensure file readability, including 2-s margins at the beginning and end of each 6-h file.

### Experiments 1 and 2

Experiment 1 consisted of divided cohabitation for 72 h starting at 8 am, i.e., 3 h after the onset of the light phase. Each male-female pair was placed in a bedded home cage (48.3 cm length, 25.4 cm width, 20.3 cm height) with a lab-made divider at the center of the cage. Thus, each animal was spatially restricted to a quadrant, producing uniform behavioral variables across recordings (**Fig 1B**). The dividers were made of metal wire mesh (square mesh, 6.5 mm aperture, 22 cm wide, 28 cm high) covered on both sides with clear polycarbonate sheet (0.5 mm thick) to prevent animals from climbing. The bottom section of the metal mesh, measuring 6.5 cm in height, was left exposed without the polycarbonate sheet, allowing animals to exchange bedding, smell each other and engage in face-to-face and body-to-body interactions. Chow and gel were placed at consistent locations away from the divider, on the left and right sides of the animal, respectively. No environmental enrichment objects were provided, encouraging the animals to interact. A handwritten label was placed outside each quadrant displaying animal metadata (sex, ELSD vs. Ctrl, quadrant number), clearly visible in each recording for extra documentation. Cardboard barriers were placed between adjacent cages, preventing different dyads from seeing each other during the multi-dyad recordings.

Experiment 2 consisted of undivided cohabitation for 4 h, also beginning at 8 am, 3-4 days after the end of Experiment 1; animals were single-housed at the vivarium between experiments. Before each recording of Experiment 2, the male and female were placed in the same home-cage quadrants, with the same handwritten label, same bedding and without chow or gel. We then removed the divider *while recording*, ensuring animal traceability throughout the recording, as male and female lacked physical differentiators (**Fig 1C**). See the next section for further identity verification methods. The male and female were then allowed to interact freely. At the end of the recording, each animal was visually inspected to confirm sex, resulting in a 100% match with the original metadata.

### Data pre-processing and animal tracking

For Experiment 1, each camera recorded two divided cages and, therefore, four quadrants, one animal per quadrant. Thus, we decided to crop each video into four smaller videos (384 x 416 pixels), one per quadrant, using Adobe Premiere and Media Encoder. By doing this we eliminated the need for identity verification in Experiment 1 (unlike Experiment 2, as described below). The cropped videos were then processed through DeepLabCut, a supervised keypoint tracker [28]. A resnet-50 network was used to label seven keypoints per animal: nose, left/right ears, three locations along the “spine”, and tail base (**Fig 1B**; **S1 Movie**). The network was trained with video frames from three males and three females, each frame representing varying levels of difficulty: from clear imaging of the animal (without obstruction of body parts) to challenging situations (e.g., curled posture when sleeping or eating, tail base hidden under bedding, nose hidden by the mesh divider). The network was trained using a laboratory server (CPU: Intel Xeon E5-2640 v3 @ 2.60 GHz. RAM memory: 512 GB. GPU: NVIDIA GeForce GTX 1080 Ti) and deployed to unseen (held-out) data. Once keypoint coordinates were generated for each recording, we proceeded to behavioral decomposition analysis using MATLAB (MathWorks), as detailed in the next section.

For Experiment 2, all video processing and behavior annotation steps were performed using LabGym2, an open-source machine vison tool that we co-developed (including authors: Y.H., B.Y., and L.S.B-J.) [32]. First, each raw video was cropped into two rectangular videos (800 x 450 pixels), one per cage, so that each video contained one male-female dyad. The cropped videos were then contrast-enhanced to highlight the animals against the background (bedded cage). These videos were then analyzed using two neural networks that we trained during our previous study: *Detector* and *Categorizer* [32]. Step 1: Segmentation and masking of each animal using the *Detector*. Step 2: Mask-based foreground extraction to obtain the raw pixels of each animal; background pixels were set to zero (black). Step 3: Unsupervised tracking of masks over consecutive frames based on foreground appearance and distance. Step 4: Generation of spatiotemporal patterns as inputs for the *Categorizer*, including body contour outlines around the animals. Specifically for the focal animal, these outlines were color-coded from blue to red to indicate the temporal progression of movements within groups of frames, i.e., time bins totaling 0.67 s per bin. The non-focal (or reference) animal was color coded in a fixed tone of gray (**Fig 1C**). The focal and reference roles were alternated per animal by processing each recording twice, enabling social role identification [32]. Time bin size (0.67 s) was determined to match the timescale of prairie vole behavior, including quick bouts of chasing and aggression. Step 5: Categorization of spatiotemporal patterns into behaviors (**Fig 1C**). This step involved manual sorting of behavioral examples – the primary supervised aspect of LabGym2 – followed by deep learning of the spatiotemporal features of these behaviors using a custom neural network structure, as detailed in our software publication [32]. LabGym2 was run remotely on the University of Michigan Great Lakes cluster. The sessions on this cluster were configured with the following resources: 2.9 GHz Intel Xeon Gold 6226R with 372 GB RAM, NVIDIA A40 with 48 GB VRAM.

LabGym2’s publication included a proof-of-concept study for prairie vole dyads, demonstrating 100% accurate individual animal tracking throughout each recording [32]. However, the recordings in that study were relatively short (10 min), whereas the recordings in the present study were longer (4 h), which we found to increase susceptibility to identity switching. Tracking identities over extended periods (e.g., several hours) indeed remains a challenging task in the field of computational ethology, despite the development of various approaches to multi-animal tracking [26–29]. In our experience, species-typical huddling behaviors exacerbated the risk of identity switching, especially 2-3 hours into the recording, when the voles tended to huddle more. Thus, we chose to identify male and female *manually*, independently on LabGym2’s behavioral categorization. Additionally, we leveraged this manual inspection effort to annotate “attacking” and “being attacked” events (**S2 Movie**), which we could not reliably automate in the current dataset. The annotation of “chasing” and “being chased”, on the other hand, was fully automated, serving as a control for our findings on sex-specific aggression roles. For the manual annotation of animal identities and aggression roles we used the graphical user interface (GUI) of another software tool: DeepEthogram [56].

In summary, we employed two distinct machine vision pipelines in this study, each tailored to a recording environment (divided and undivided cohabitation). However, our study also involved significant human supervision, including manual validation of sex identities and aggression roles, mitigating the risk of over-automation [57]. In fact, our results on “chasing” and “attacking” supported each other (**Fig 3**), which is an important control, as “chasing” and “attacking” were annotated by computer and human, respectively. We believe that human-in-the-loop strategies like these will continue to offer a balance between effort cost and ethological validity, revealing meaningful patterns like the neurotype matching effects reported here.

### Behavioral decomposition and categorization

<u>Experiment 1</u>: Keypoints from DeepLabCut represented each animal as an arrow-shaped “skeleton”, one per video frame (**Fig 1B**). The coordinates for these keypoints were imported into MATLAB, converted from pixels to centimeters, and processed into three behavioral variables.

1. Direction to divider: a line was fitted between the extreme keypoints (nose and tail base), and the angle of this line relative to the horizontal plane of the frame was calculated. This angle was then rescaled from −1 (opposite from the divider) to +1 (toward the divider), with clockwise and counterclockwise directions treated equally.
2. Distance to divider: horizontal distance between the center of the animal and the divider, regardless of its position along the vertical axis.
3. Locomotion speed: difference (hypotenuse) between the center of the animal in frame n and frame n+1.

These measures were timestamped according to clock time, in number of seconds x number of frames. The clock was not restarted at midnight, assigning each frame to a unique temporal identifier across the 72 hours. This allowed us to trim and align the recordings to a common 72 h axis, as safety margins of a few minutes were recorded before and after each session. Clock time information was obtained from file naming metadata through the acquisition software (StreamPix, NorPix). These procedures replicate those used in a previous publication [31]. See **S1 Movie**.

<u>Experiment 2</u>: LabGym2’s outputs include the likelihood of each behavioral category per video frame on a 0-1 scale. The categories were initially the same as our proof-of-concept study in prairie voles [32]: aggression, chasing, being chased, approaching, sniffing, being sniffed, moving alone, idling alone, huddling. For the present study, aggression was ignored (replaced by manual annotations, as described below) and adaptations to other categories were made using MATLAB.

1. Sniffing into Pointing: This category merged three types of sniffing (face-to-face, face-to-body, anogenital) and approaching behaviors. This was renamed to a generic term, “pointing”, as freezing-like behaviors that typically occurred after aggressive bouts (**S2 Movie**) were often misclassified as “sniffing” under LabGym2’s spatiotemporal patterning. Neural network refinement and deeper ethological exploration were outside our scope (e.g., sub-classification of sniffing, sequencing of peri-aggression postures).
2. Behaving alone: This category merged “moving alone” and “idling alone” (this latter included self-grooming). In-depth exploration of these solitary behaviors was outside our scope, hence the creation of this generic category. Frames were set to zero likelihood of “behaving alone” if only one animal was classified under this category, making it a full-dyad category (see purple curves in **Fig 3**).
3. Huddling: Same as the previous proof-of-concept study in 10-min recordings [32], but with one correction that we found necessary in the present 4-h recordings. During intense body contact, the identities of both animals sometimes merged into one, causing “huddling” to be misclassified as “idling alone”. Frames with this issue we re-classified as huddling. Frames were set to zero likelihood of “huddling” if only one animal was classified under this category, making it a full-dyad category (see purple curves in **Fig 3**).

After these adaptations, likelihood values were converted into a vector of integer labels using a winner-takes-all approach. Finally, frames manually classified as “attacking” and “being attacked” were superimposed on that vector of labels. For final quantification (explained next), the focal roles of chasing, attacking, and pointing were analyzed without their mirroring counterparts (i.e., being chased, being attacked, being pointed at), thus minimizing redundancy in the figures.

### Final data processing and statistical analysis per figure

**Figs 2 and 3**: Time series data obtained from the behavioral decomposition (Experiment 1) and behavioral annotation (Experiment 2) methods above were analyzed, with each data point corresponding to one video frame of a single animal. These data points were binned and smoothed on a per-animal basis: the 72-h recordings of Experiment 1 were averaged into 1-h bins (total of 72 bins) and the 4-h recordings of Experiment 2 were averaged into 3-min bins (total of 80 bins), both followed by a 6-bin moving mean window. The binned data were then plotted as +/- standard error curves comparing males vs. females overall, males vs. females within major dyad groups (matched or mixed), males vs. females within dyad sub-groups (Ctrl-Ctrl, ELSD-ELSD, Ctrl-ELSD, ELSD-Ctrl), and matched vs. mixed dyads within each sex. These comparisons were performed comprehensively across behavioral variables, allowing to identify sex or dyad type differences with both: (1) behavioral variable specificity (in which behavioral variables the dyad type precipitated or amplified differences?); and (2) temporal specificity (at which time bins did these differences emerge?). Statistical comparisons were made using two-way ANOVA with time bins as repeated measures, followed by Tukey’s post-hoc tests per time bin. The main statistical effects (sex, time, interaction) are reported in **Tables 1 and 2** and significant post-hoc differences are depicted as bold lines along the x-axes in **Figs 2 and 3**. Distributions per repeated measure passed the Kolmogorov-Smirnov normality test.

**Fig 4**: Experiments 1 and 2 were performed in different environments (divided and undivided cohabitation), 3-4 days apart. The animals and dyad assignments were the same between experiments, allowing for correlation analyses among all eight variables from the two experiments. To investigate this, behavioral variables were averaged across the entirety of each experiment, resulting in eight averages per animal. These averages were then grouped by sex and major dyad type (males or females from matched or mixed), forming four animal x variable arrays. Linear correlations within each sex and dyad type group were calculated, producing matrices with correlation coefficients (R) and corresponding P values. Key correlations were illustrated as scatter plots with linear fits in **Fig 4A-B**, while the R and P value matrices themselves were illustrated with a color code in **Fig 4C**. Distributions per variable passed the Kolmogorov-Smirnov normality test.

**Fig 5A-B**: Behavior transition probabilities were computed using 1-s time bins, matching the approximate timescale of prairie vole behaviors, including chasing and aggression bouts. For Experiment 1, continuous DeepLabCut-based variables of direction and distance to divider were averaged per time bin and discretized into categorical vectors (number of elements = number of time bins), enabling transition probability analysis. For discretization, direction to divider was converted from a −1 to 1 scale into “toward” vs. “opposite”, whereas distance to divider was converted from a centimeter scale into “near” vs. “away”. These categories were combined into a four-category vector (toward-near, opposite-near, toward-away, opposite-away), one per animal. Then each dyad had a “time t” animal and a “time t+1” animal assignment, one iteration per sex directionality (male-then-female, female-then-male). Each of these iterations resulted in a 4 x 4 adjacency matrix (4 = number of behavioral categories), with rows representing “time t” behaviors, columns representing “time t+1” behaviors, and values indicating probabilities. Multiple such adjacency matrices were created, grouped by both sex directionality and dyad type. Specifically for the illustrations in **Fig 5A**, adjacency matrices were averaged across the dyads of each group and plotted as directed graphs. The edges of these graphs were coded by both weight and grayscale: the thicker and darker the edge, the higher the transition probability. For quantification (**Fig 5B**) no averaging was applied. Instead, each adjacency matrix (one per dyad) was reshaped into a row vector containing all behavior pairs (“behavior in time t” and “behavior in time t+1”). These rows were then concatenated across dyads, creating dyad x behavior pair matrices. Finally, dyad x behavior pair matrices from matched and mixed dyads were statistically compared per sex directionality using two-way ANOVA with behavior pairs as repeated measures, followed by Tukey’s post-hoc tests per behavior pair. The main statistical effects (dyad type, behavior pair, interaction) are reported in **Table 3** and significant post-hoc differences are depicted as bold edges in **Fig 5B**. Distributions per repeated measure passed the Kolmogorov-Smirnov normality test.

**Fig 5C-D**: Similar methods were used for Experiment 2 with three differences: (1) LabGym2-based data were already in categorical form prior to analysis, eliminating the need for categorization; (2) each time bin (1 s) was assigned its most frequent category (winner-takes-all); and (3) this analysis included five behavioral categories (chasing, attacking, pointing, behaving alone, huddling), leading to 5 x 5 adjacency matrices.

## Data and code availability

The behavior tracking software (LabGym) and the training dataset for prairie voles are both available on GitHub (https://github.com/umyelab/LabGym). Processed post-tracking data, as well as code for figure plotting and statistical analysis are available to peer reviewers via private link on figshare.

## Supporting information

**S1 Movie.** Decomposing behavioral variables in the divided cage

**S2 Movie.** Examples of behaviors in the undivided cage

## Acknowledgements

This research was supported by the National Institute of Mental Health to B.O.W., M.M.L., L.S.B-J., C.E.J-T. (R01MH131592) and N.E.P.M. (F31MH136684), and the National Institute of Neurological Disorders and Stroke to B.Y. (R01NS137222). We thank all lab colleagues for discussions.

## Author contributions

**Conceptualization**: Lezio S. Bueno-Junior, Noah E. P. Milman, Carolyn E. Jones-Tinsley, Miranda M. Lim, Brendon O. Watson.

**Data curation**: Lezio S. Bueno-Junior, Brendon O. Watson.

**Formal analysis**: Lezio S. Bueno-Junior.

**Funding acquisition**: Brendon O. Watson, Miranda M. Lim, Lezio S. Bueno-Junior, Carolyn E. Jones-Tinsley.

**Investigation**: Lezio S. Bueno-Junior.

**Methodology**: Lezio S. Bueno-Junior, Carolyn E. Jones-Tinsley, Noah E. P. Milman.

**Project administration**: Brendon O. Watson, Miranda M. Lim.

**Software**: Lezio S. Bueno-Junior, Yujia Hu, Bing Ye, Anjesh Ghimire.

**Resources**: Carolyn E. Jones-Tinsley, Peyton T. Wickham, Brendon O. Watson, Miranda M. Lim.

**Supervision**: Brendon O. Watson, Miranda M. Lim.

**Visualization**: Lezio S. Bueno-Junior.

**Writing – original draft**: Lezio S. Bueno-Junior.

**Writing – review & editing**: Noah E. P. Milman, Brendon O. Watson, Miranda M. Lim, Carolyn E. Jones-Tinsley, Anjesh Ghimire, Peyton T. Wickham, Yujia Hu, Bing Ye.

