## Supplementary material for "Neurotype matching in monogamous rodents is modulated by early-life sleep experience": Table 1

| figure panel | dyad type or sex | effect | value | direction to divider | distance to divider | locomotion speed |
|---|---|---|---|---|---|---|
| 2A | all dyads | sex | deg fr | 1, 27 | 1, 27 | 1, 27 |
| | | | F; P | **=37.78; <0.001** | =0.13; =0.722 | =4.03; =0.055 |
| | | time | deg fr | 71, 1917 | 71, 1917 | 71, 1917 |
| | | | F; P | =7.92; <0.001 | =9.06; <0.001 | =19.10; <0.001 |
| | | interaction | deg fr | 71, 1917 | 71, 1917 | 71, 1917 |
| | | | F; P | **=3.13; <0.001** | =1.01; =0.460 | **=1.73; <0.001** |
| 2B | matched dyads | sex | deg fr | 1, 14 | 1, 14 | 1, 14 |
| | | | F; P | =7.72; =0.015 | =1.50; =0.241 | =0.85; =0.373 |
| | | time | deg fr | 71, 994 | 71, 994 | 71, 994 |
| | | | F; P | =4.46; <0.001 | =3.51; <0.001 | =11.10; <0.001 |
| | | interaction | deg fr | 71, 994 | 71, 994 | 71, 994 |
| | | | F; P | =1.06; =0.340 | =0.73; =0.955 | =0.76; =0.929 |
| | mixed dyads | sex | deg fr | 1, 12 | 1, 12 | 1, 12 |
| | | | F; P | **=72.53; <0.001** | =1.30; =0.276 | =4.52; =0.055 |
| | | time | deg fr | 71, 852 | 71, 852 | 71, 852 |
| | | | F; P | =4.33; <0.001 | =6.06; <0.001 | =8.88; <0.001 |
| | | interaction | deg fr | 71, 852 | 71, 852 | 71, 852 |
| | | | F; P | **=4.52; <0.001** | =1.05; =0.373 | **=1.63; =0.001** |
| 2C | Ctrl-Ctrl | sex | deg fr | 1, 6 | 1, 6 | 1, 6 |
| | | | F; P | =3.04; =0.132 | =0.39; =0.553 | =0.15; =0.715 |
| | | time | deg fr | 71, 426 | 71, 426 | 71, 426 |
| | | | F; P | =3.54; <0.001 | =2.80; <0.001 | =5.74; <0.001 |
| | | interaction | deg fr | 71, 426 | 71, 426 | 71, 426 |
| | | | F; P | =0.77; =0.915 | =0.49; =1.000 | =0.41; =1.000 |
| | ELSD-ELSD | sex | deg fr | 1, 7 | 1, 7 | 1, 7 |
| | | | F; P | =4.27; =0.078 | =1.17; =0.315 | =1.71; =0.232 |
| | | time | deg fr | 71, 497 | 71, 497 | 71, 497 |
| | | | F; P | =2.14; <0.001 | =1.51; =0.007 | =5.39; <0.001 |
| | | interaction | deg fr | 71, 497 | 71, 497 | 71, 497 |
| | | | F; P | =0.89; =0.720 | =0.61; =0.995 | =1.00; =0.485 |
| | Ctrl-ELSD | sex | deg fr | 1, 6 | 1, 6 | 1, 6 |
| | | | F; P | **=35.70; <0.001** | =5.02; =0.066 | =1.10; =0.335 |
| | | time | deg fr | 71, 426 | 71, 426 | 71, 426 |
| | | | F; P | =3.86; <0.001 | =3.78; <0.001 | =3.13; <0.001 |
| | | interaction | deg fr | 71, 426 | 71, 426 | 71, 426 |
| | | | F; P | **=2.31; <0.001** | **=2.57; <0.001** | =0.41; =1.000 |
| | ELSD-Ctrl | sex | deg fr | 1, 5 | 1, 5 | 1, 5 |
| | | | F; P | **=37.71; =0.002** | =0.05; =0.838 | =3.62; =0.115 |
| | | time | deg fr | 71, 355 | 71, 355 | 71, 355 |
| | | | F; P | =1.24; =0.111 | =2.77; <0.001 | =7.65; <0.001 |
| | | interaction | deg fr | 71, 355 | 71, 355 | 71, 355 |
| | | | F; P | **=2.52; <0.001** | =0.35; =1.000 | **=3.46; <0.001** |
| 2D | males | dyad type | deg fr | 1, 12 | 1, 12 | 1, 12 |
| | | | F; P | =8.26; =0.014 | =0.96; =0.346 | =0.03; =0.865 |
| | | time | deg fr | 71, 852 | 71, 852 | 71, 852 |
| | | | F; P | =3.15; <0.001 | =3.44; <0.001 | =8.36; <0.001 |
| | | interaction | deg fr | 71, 852 | 71, 852 | 71, 852 |
| | | | F; P | **=1.58; =0.002** | =0.54; =0.999 | =0.55; =0.999 |
| | females | dyad type | deg fr | 1, 12 | 1, 12 | 1, 12 |
| | | | F; P | =1.42; =0.257 | =2.05; =0.178 | =0.28; =0.607 |
| | | time | deg fr | 71, 852 | 71, 852 | 71, 852 |
| | | | F; P | =7.47; <0.001 | =7.12; <0.001 | =9.06; <0.001 |
| | | interaction | deg fr | 71, 852 | 71, 852 | 71, 852 |
| | | | F; P | =1.09; =0.302 | =0.97; =0.540 | =1.30; =0.055 |

**Table 1. Statistics for Figure 2.** Degrees of freedom, F, and P values were obtained using two-way ANOVA with time bins as repeated measures. P values < 0.01 were highlighted with bold font, except for repeated measure effects.
