## Supplementary material for "Neurotype matching in monogamous rodents is modulated by early-life sleep experience": Table 2

| figure panel | dyad type or sex | effect | value | chasing | attacking | pointing | behaving alone | huddling |
|---|---|---|---|---|---|---|---|---|
| 3A | all dyads | sex | deg fr | 1, 27 | 1, 27 | 1, 27 | 1, 27 | 1, 27 |
| | | | F; P | =4.79; =0.038 | **=16.28; <0.001** | =1.11; =0.302 | =0.00; =1.000 | =0.00; =1.000 |
| | | time | deg fr | 79, 2133 | 79, 2133 | 79, 2133 | 79, 2133 | 79, 2133 |
| | | | F; P | =19.20; <0.001 | =21.83; <0.001 | =8.95; <0.001 | =8.64; <0.001 | =13.89; <0.001 |
| | | interaction | deg fr | 79, 2133 | 79, 2133 | 79, 2133 | 79, 2133 | 79, 2133 |
| | | | F; P | **=3.89; <0.001** | **=11.30; <0.001** | =0.95; =0.607 | =0.00; =1.000 | =0.00; =1.000 |
| 3B | matched dyads | sex | deg fr | 1, 14 | 1, 14 | 1, 14 | 1, 14 | 1, 14 |
| | | | F; P | =0.31; =0.588 | =3.77; =0.073 | =0.72; =0.409 | =0.00; =1.000 | =0.00; =1.000 |
| | | time | deg fr | 79, 1106 | 79, 1106 | 79, 1106 | 79, 1106 | 79, 1106 |
| | | | F; P | =8.72; <0.001 | =12.96; <0.001 | =4.54; <0.001 | =6.06; <0.001 | =8.66; <0.001 |
| | | interaction | deg fr | 79, 1106 | 79, 1106 | 79, 1106 | 79, 1106 | 79, 1106 |
| | | | F; P | =0.61; =0.997 | **=4.25; <0.001** | =0.64; =0.993 | =0.00; =1.000 | =0.00; =1.000 |
| | mixed dyads | sex | deg fr | 1, 12 | 1, 12 | 1, 12 | 1, 12 | 1, 12 |
| | | | F; P | =6.87; =0.022 | **=15.96; =0.002** | =0.35; =0.565 | =0.00; =1.000 | =0.00; =1.000 |
| | | time | deg fr | 79, 948 | 79, 948 | 79, 948 | 79, 948 | 79, 948 |
| | | | F; P | =12.11; <0.001 | =13.83; <0.001 | =4.77; <0.001 | =3.08; <0.001 | =5.38; <0.001 |
| | | interaction | deg fr | 79, 948 | 79, 948 | 79, 948 | 79, 948 | 79, 948 |
| | | | F; P | **=5.25; <0.001** | **=8.37; <0.001** | =0.88; =0.757 | =0.00; =1.000 | =0.00; =1.000 |
| 3C | Ctrl-Ctrl | sex | deg fr | 1, 6 | 1, 6 | 1, 6 | 1, 6 | 1, 6 |
| | | | F; P | =0.06; =0.822 | =8.63; =0.026 | =0.35; =0.578 | =0.00; =1.000 | =0.00; =1.000 |
| | | time | deg fr | 79, 474 | 79, 474 | 79, 474 | 79, 474 | 79, 474 |
| | | | F; P | =4.49; <0.001 | =3.82; <0.001 | =2.33; <0.001 | =1.42; =0.015 | =2.75; <0.001 |
| | | interaction | deg fr | 79, 474 | 79, 474 | 79, 474 | 79, 474 | 79, 474 |
| | | | F; P | =0.52; =1.000 | **=3.58; <0.001** | =0.29; =1.000 | =0.00; =1.000 | =0.00; =1.000 |
| | ELSD-ELSD | sex | deg fr | 1, 7 | 1, 7 | 1, 7 | 1, 7 | 1, 7 |
| | | | F; P | =0.91; =0.373 | =0.31; =0.598 | =3.47; =0.105 | =0.00; =1.000 | =0.00; =1.000 |
| | | time | deg fr | 79, 553 | 79, 553 | 79, 553 | 79, 553 | 79, 553 |
| | | | F; P | =6.16; <0.001 | =9.99; <0.001 | =2.48; <0.001 | =6.15; <0.001 | =6.72; <0.001 |
| | | interaction | deg fr | 79, 553 | 79, 553 | 79, 553 | 79, 553 | 79, 553 |
| | | | F; P | =0.68; =0.981 | **=1.45; =0.010** | **=1.76; <0.001** | =0.00; =1.000 | =0.00; =1.000 |
| | Ctrl-ELSD | sex | deg fr | 1, 6 | 1, 6 | 1, 6 | 1, 6 | 1, 6 |
| | | | F; P | =3.54; =0.109 | =9.11; =0.023 | =0.67; =0.444 | =0.00; =1.000 | =0.00; =1.000 |
| | | time | deg fr | 79, 474 | 79, 474 | 79, 474 | 79, 474 | 79, 474 |
| | | | F; P | =6.24; <0.001 | =6.58; <0.001 | =3.84; <0.001 | =2.30; <0.001 | =3.53; <0.001 |
| | | interaction | deg fr | 79, 474 | 79, 474 | 79, 474 | 79, 474 | 79, 474 |
| | | | F; P | **=3.86; <0.001** | **=6.76; <0.001** | =0.65; =0.990 | =0.00; =1.000 | =0.00; =1.000 |
| | ELSD-Ctrl | sex | deg fr | 1, 5 | 1, 5 | 1, 5 | 1, 5 | 1, 5 |
| | | | F; P | =2.92; =0.148 | =5.97; =0.058 | =0.06; =0.816 | =0.00; =1.000 | =0.00; =1.000 |
| | | time | deg fr | 79, 395 | 79, 395 | 79, 395 | 79, 395 | 79, 395 |
| | | | F; P | =5.91; <0.001 | =7.17; <0.001 | =1.96; <0.001 | =1.91; <0.001 | =2.85; <0.001 |
| | | interaction | deg fr | 79, 395 | 79, 395 | 79, 395 | 79, 395 | 79, 395 |
| | | | F; P | **=2.20; <0.001** | **=3.28; <0.001** | =0.57; =0.999 | =0.00; =1.000 | =0.00; =1.000 |
| 3D | males | dyad type | deg fr | 1, 12 | 1, 12 | 1, 12 | 1, 12 | 1, 12 |
| | | | F; P | =0.60; =0.454 | =0.01; =0.943 | =1.72; =0.215 | =0.67; =0.430 | =0.11; =0.746 |
| | | time | deg fr | 79, 948 | 79, 948 | 79, 948 | 79, 948 | 79, 948 |
| | | | F; P | =11.40; <0.001 | =6.25; <0.001 | =5.16; <0.001 | =6.94; <0.001 | =11.35; <0.001 |
| | | interaction | deg fr | 79, 948 | 79, 948 | 79, 948 | 79, 948 | 79, 948 |
| | | | F; P | =0.90; =0.716 | =0.82; =0.873 | =0.66; =0.990 | =0.45; =1.000 | =0.35; =1.000 |
| | females | dyad type | deg fr | 1, 12 | 1, 12 | 1, 12 | 1, 12 | 1, 12 |
| | | | F; P | **=10.81; =0.006** | =5.82; =0.033 | =2.77; =0.122 | =0.67; =0.430 | =0.11; =0.746 |
| | | time | deg fr | 79, 948 | 79, 948 | 79, 948 | 79, 948 | 79, 948 |
| | | | F; P | =13.05; <0.001 | =19.21; <0.001 | =8.67; <0.001 | =6.935; <0.001 | =11.35; <0.001 |
| | | interaction | deg fr | 79, 948 | 79, 948 | 79, 948 | 79, 948 | 79, 948 |
| | | | F; P | **=2.70; <0.001** | **=4.42; <0.001** | =0.36; =1.000 | =0.448; =1.000 | 0.35; =1.000 |

**Table 2. Statistics for Figure 3.** Degrees of freedom, F, and P values were obtained using two-way ANOVA with time bins as repeated measures. P values < 0.01 were highlighted with bold font, except for repeated measure effects.
