## Supplementary material for "Neurotype matching in monogamous rodents is modulated by early-life sleep experience": Table 3

| figure panel | sex direction | effect | value | |
|---|---|---|---|---|
| 5B | male-to-female | dyad type | deg fr | 1, 12 |
| | | | F; P | =0.02; =0.904 |
| | | behavior pair | deg fr | 11, 132 |
| | | | F; P | =123.86; <0.001 |
| | | interaction | deg fr | 11, 132 |
| | | | F; P | =0.69; =0.747 |
| | female-to-male | dyad type | deg fr | 1, 12 |
| | | | F; P | =0.01; =0.955 |
| | | behavior pair | deg fr | 11, 132 |
| | | | F; P | =138.55; <0.001 |
| | | interaction | deg fr | 11, 132 |
| | | | F; P | **=5.23; <0.001** |
| 5D | male-to-female | dyad type | deg fr | 1, 12 |
| | | | F; P | =0.80; =0.389 |
| | | behavior pair | deg fr | 19, 228 |
| | | | F; P | =70.03; <0.001 |
| | | interaction | deg fr | 19, 228 |
| | | | F; P | =0.87; =0.618 |
| | female-to-male | dyad type | deg fr | 1, 12 |
| | | | F; P | =5.13; =0.043 |
| | | behavior pair | deg fr | 19, 228 |
| | | | F; P | =68.26; <0.001 |
| | | interaction | deg fr | 19, 228 |
| | | | F; P | **=2.64; <0.001** |

**Table 3. Statistics for Figure 5.** Degrees of freedom, F, and P values were obtained using two-way ANOVA with behavior pairs as repeated measures. P values < 0.01 were highlighted with bold font, except for repeated measure effects.
